# Synaptic Alterations Are Preceding the Axonal Loss in Optic Atrophy of Wolfram Syndrome Mouse Model

**DOI:** 10.64898/2026.03.22.713521

**Authors:** Venu Gurram, William An, Shrinivas Bimal, Fumihiko Urano

## Abstract

Wolfram syndrome is a rare autosomal recessive disorder characterized by antibody-negative early-onset diabetes mellitus, optic atrophy, sensorineural hearing loss, arginine-vasopressin deficiency, and progressive neurodegeneration of the brainstem and cerebellum. It is caused primarily by pathogenic variants in the *WFS1* gene, which encodes a transmembrane endoplasmic reticulum-resident protein involved in the unfolded protein response and cellular calcium homeostasis. Although multiple rodent models of Wolfram syndrome have been developed and shown to exhibit visual defects, some studies have reported significant vision loss prior to any detectable axonal degeneration or myelin abnormalities, and the mechanisms underlying these early visual deficits remain poorly understood. Recent *in vitro* studies have demonstrated altered synaptic contacts and aberrant neurite morphology in *WFS1*-deficient cerebral organoids and human iPSC-derived neurons, respectively. These findings prompted us to investigate, for the first time in vivo, whether synaptic and dendritic abnormalities occur in the retina of *Wfs1* knockout mice. Using confocal microscopy, we examined retinal and optic nerve histology in *Wfs1* knockout mice at 4 and 7 months of age. Our analysis reveals progressive synaptic alterations in the inner plexiform layer, driven by early presynaptic compartment failure. These changes represent the earliest detectable phenotype associated with vision loss in this model and precede overt axonal degeneration.

## 1. Introduction

Wolfram syndrome is an rare life-threatening neurodegenerative disorder, with no treatment, inherited in autosomal recessive manner affecting multiple organs resulting in early-onset antibody-negative diabetes mellitus, optic atrophy, sensorineural hearing loss, arginine-vasopressin deficiency, and cerebellar and brainstem degeneration with a prevalence of 1 in 160,000–770,000 [1; 2; 3; 4]. The life expectancy of the majority of patients ranges from 30–40 years and respiratory failure or dysphagia are the common causes of death [4; 5].

Wolfram syndrome is mainly caused by homozygous or compound heterozygous pathogenic variant in the WFS1 gene encoding a transmembrane protein localized to the endoplasmic reticulum (ER), wolframin [6]. Wolframin is an 890-amino acid protein with a molecular mass of 100 kDa with its C-terminus oriented towards the ER lumen and its N-terminus towards the cytoplasm, and it can assemble and form tetramers [6; 7; 8]. WFS1 has been shown to play roles in the unfolded protein response, cellular calcium regulation, mitochondrial-associated membranes, and ER vesicular cargo export. [9; 10; 11; 12; 13; 14; 15; 16].

Optic atrophy is of serious concern in patients with Wolfram syndrome due to its progressive nature. Case reports on patients with Wolfram syndrome have revealed progressive loss of visual acuity, color vision, and cecocentral scotomas starting in their early second decade of life, indicating the retinal ganglion cells (RGCs) as the primary pathological site [17; 18; 19; 20]. Several research groups have studied optic atrophy in Wolfram syndrome using animal models. One study with the *Wfs1*-knockout model reported abnormal retinal function and slower conduction along the visual pathways with no RGC loss but with reduced axonal density compared to wildtype controls at 12 months of age [21]. Another study showed a significant reduction in the ratio of retinal thickness to longitudinal diameter and a decreased number of GFAP-positive cells in the inner nuclear layer (INL) as well as reduced GFAP intensity in the retina of *Wfs1*-knockout mice compared to control mice at approximately 3.5 months of age [22]. Recently, a study reported visual impairments including reduced visual acuity, axonal loss, myelin degeneration, increased retinal gliosis, but no RGC loss at 12 months of age in *Wfs1*-knockout mice [23]. In another study, progressive visual defects were reported starting as early as 3 months of age and myelin defects at 7.5 months in Wfs1 knockout mice [24]. A *Wfs1*-knockout rat model showed reduced visual acuity starting at 8.5 months, and axonal loss and increased retinal gliosis at 14–15 months, with significant RGC loss at 17–18 months [25; 26; 27]. In a zebrafish *wfs1*-knockout model, significant RGC loss was observed at 4 months and a thinner GCL at 12 months, with significantly reduced visual acuity at both ages [28].

A recent study showed disrupted synapses and altered neurite outgrowth in *Wfs1*-deficient cerebral organoids compared to control organoids [29]. Additionally, altered neurite outgrowth was also observed in human iPSC-derived neurons deficient for *Wfs1* [30]. It is therefore necessary to investigate synaptic and dendritic alterations in vivo in Wolfram syndrome model. Although multiple previous studies have investigated optic atrophy in Wolfram syndrome, none have focused on synaptic and dendritic arborization in the retina of a Wolfram syndromemouse model. Therefore, this study aims to provide comprehensive, age-dependent phenotyping of RGCs by examining their cell bodies in the retina and their axons in the optic nerve in a *Wfs1*-knockout mouse model of Wolfram syndrome.

## 2. Materials and Methods

### 2.1 Animals

The 129S6 mice lacking *Wfs1* expression throughout the body were originally provided by Dr. Sulev Kõks at the University of Tartu [31; 32]. In this strain, the segment of the WFS1 protein spanning amino acids 360 to 890 was substituted with an NLS LacZ Neo cassette inserted in frame. Genotyping was carried out using multiplex PCR performed by Transnetyx (Cordova, TN). All procedures involving animals followed the guidelines approved by the Washington University School of Medicine IACUC (Protocol #20 0334). Mice were maintained in a barrier-protected facility with continuous, unrestricted access to food and water for the duration of the study.

### 2.2 Tissue Collection and Preparation

Mice were euthanized in a CO chamber for 7 minutes, and eyes and optic nerves were collected immediately. For RNA and protein analysis, tissues were flash-frozen and stored at −80°C until use. For histology, optic nerves and eye cups (following removal of the cornea and lens) were fixed overnight at 4°C in 4% paraformaldehyde. Tissues were then cryoprotected in 30% sucrose in PBS at 4°C overnight. Eyes were subsequently embedded in OCT and stored at −20°C. Optic nerves were embedded in gelatin as previously described [33] and stored at −80°C.

### 2.3 Immunohistochemistry and Imaging

Retinas and optic nerves were sectioned at 10 μm using a cryostat (Leica CM1950) and mounted on SuperFrost Plus slides (12-550-15, Fisherbrand). Following PBS rinsing, sections were permeabilized with 0.25% Triton X-100 for 20 min at room temperature and blocked for 2 h in PBS containing 3% BSA, 5% goat serum, and 0.1% Triton X-100. Primary antibodies against Brn3a (MAB1585, Millipore, 1:250), RBPMS (GTX118619, GeneTex, 1:200), NF200 (N4142, Sigma-Aldrich, 1:2000), GFAP (PA1-10004, Invitrogen, 1:2000), MBP aa82–87 (MCA409S, Bio-Rad, 1:50), β-III Tubulin (ab18207, Abcam, 1:750), PSD95 (51-6900, Invitrogen, 1:150), and Synaptophysin (101 004, Synaptic Systems, 1:2000) were applied overnight at 4°C. Secondary antibodies (1:1000) and DAPI (0.5 μg/mL) were applied for 1 h at room temperature. For MBP staining, optic nerve sections were incubated with ice-cold methanol for 7 min at −20°C prior to the permeabilization step.

For RGC quantification, retinal sections labeled with anti-RBPMS and anti-Brn3a antibodies were imaged using tile scan on a Leica widefield microscope at 20× to capture complete retinal sections. Images were acquired from middle sections of the retina, avoiding peripheral sections. Cell quantification was performed using the Fiji cell counter plugin. The analyst was blinded to genotype during image acquisition and analysis. Three to four retinal sections per mouse were evaluated for each genotype.

Imaging of retinal sections was performed on a Nikon AXR confocal laser-scanning microscope using a 60× objective with a z-step size of 0.3 μm and a zoom factor of 2.0. Consistent imaging settings were maintained across all samples within each experiment. On average, 12 images were acquired across a minimum of 3 retinal sections per mouse. Only middle retinal sections were imaged, focusing on the central retina and excluding peripheral regions and the optic nerve head.

Imaging of optic nerve sections was performed on Zeiss LSM 880 Airyscan confocal microscope using tile scan (2×2) with a 40× objective with z-step size of 0.22 μm and a zoom factor of 1.8. Identical microscope settings were maintained throughout all imaging sessions. A minimum of 3 images from 3 optic nerve sections per mouse were acquired to ensure adequate tissue representation.

### 2.4 Imaris Image Analysis

Image analysis was performed using Imaris (version 10.2 or later) (Bitplane). Prior to quantification, all images underwent background subtraction followed by application of a consistent threshold cutoff. These preprocessing steps were applied uniformly across all images within each experiment to ensure comparability. Depending on the analysis type, either spots or surfaces were generated in Imaris using identical parameter settings for all samples.

For NF200-positive axonal counts, the Imaris Spots module was used to create and automatically quantify spot objects. For measurements of fluorescence intensity and volume, the Surface module was used to generate 3D surface reconstructions. Colocalization analyses were performed using the surface–surface colocalization function within Imaris.

### 2.5 Immunoblot

Total protein lysates were isolated from retinal tissues using T-PER Tissue Protein Extraction Reagent (78510, ThermoFisher Scientific) supplemented with 1× complete protease inhibitor cocktail (11873580001, MilliporeSigma). Protein lysates were mixed with 4× Laemmli buffer (1610747, Bio-Rad) and heated at 45°C for 25 min. Proteins were separated by SDS-PAGE and transferred to a Nitrocellulose Membrane (pore size 0.2 μm; 1620112, Bio-Rad). Primary antibodies used were anti-WFS1 (1158-1-AP, Proteintech, 1:1000) and β-actin (4967, Cell Signaling Technology, 1:3000). Secondary antibodies conjugated to horseradish peroxidase were used for detection. Protein bands were developed with ECL Select (RPN2235, MilliporeSigma) and imaged on a ChemiDoc MP Imaging System (Bio-Rad).

### 2.6 Quantitative Real-Time PCR

Total RNA was extracted from retinal tissue using the RNeasy Mini Kit (74106, Qiagen) and reverse-transcribed using the SuperScript III First-Strand Synthesis System (18080051, Invitrogen). The resulting cDNA was analyzed by qPCR in 3 technical replicates. Primer sequences used were: *Wfs1* (Forward: 5′-CCAGCTGAGGAACTTCAAGG-3′; Reverse: 5′-AGGATGACCACGGACAGTTC-3′) and *Gapdh* (Forward: 5′-TGTAGACCATGTAGTTGAGGTCA-3′; Reverse: 5′-AGGTCGGTGTGAACGGATTTG-3′).

### 2.7 Statistical Analysis

All data were analyzed using GraphPad Prism. Data are presented as mean ± SEM relative to wild-type values unless otherwise noted. Comparisons between *Wfs1* knockout and WT mice were conducted using unpaired Student’s t-tests with Welch’s correction or Mann–Whitney tests, based on the Kolmogorov-Smirnov normality test results. The specific statistical test and p value for each individual result are detailed in the corresponding figure legend.

## 3. Results

### 3.1 No Expression of WFS1 protein and Reduced Body Weight in Wfs1 KO Mice

*Wfs1* KO mice were generated using a *Wfs1*-exon8 deletion, resulting in complete knockout of WFS1 [31; 32]. To confirm WFS1 deletion, western blot analysis was performed using whole retinal protein extract from 7-month-old animals. As expected, immunoblots showed no band in *Wfs1* knockout mice whereas a clear band at 100 kDa was detected in wildtype controls (Figure 1A). Absence of expression was confirmed by qPCR (Figure 1B). At 4 months of age, male *Wfs1* knockout mice showed a statistically significant reduction in body weight compared to male wildtype littermates (Figure 1C).

**Figure 1.**
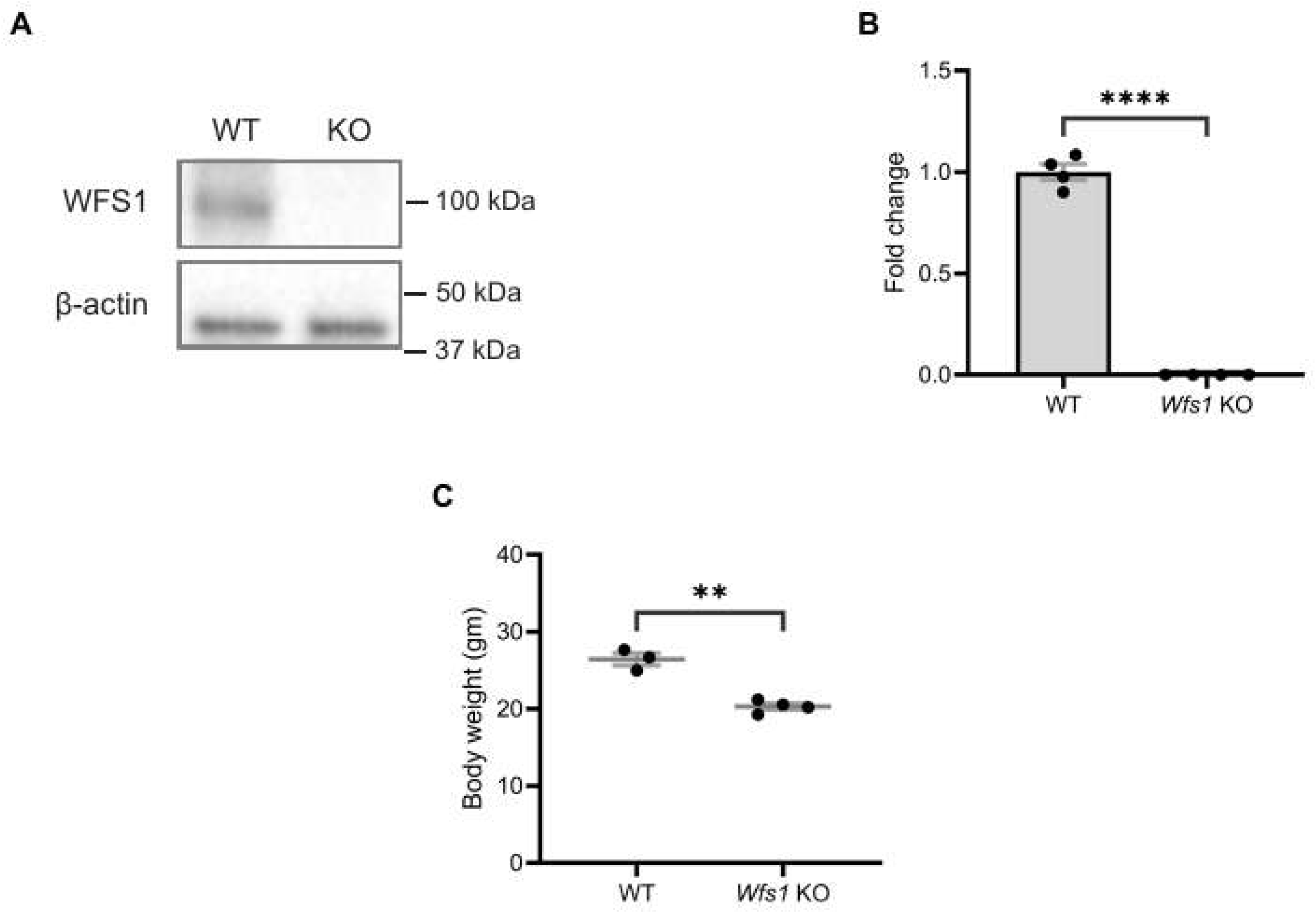
No wolframin expression and reduced body weight in *Wfs1* KO mice. (A) Representative immunoblot showing WFS1 expression in total protein lysate from retina of both wild type and *Wfs1* KO mice. β-actin was used as loading control. (8) Quantification of mRNA expression of *Wfs1* from total RNA from retina of wild type and *Wfs1* KO mice. Data are presented as normalized values ± SEM relative to WT mean. Statistical analysis was performed by two-tailed, unpaired student’s I-test with Welch’s correction. n=4. \*\*\**P*<0.0001. *Gapdh* as housekeeping gene. (C) Quantification of body weight of *Wfs1* KO mice compared to the wild type control mice at the age of 4 months. Datapoints were presented as mean ± SEM. Statistical analysis was performed by two-tailed, unpaired student’s I-test with Welch’s correction. n ≥ 3. \*\**P*<0.01.

### 3.2 No RGC Loss in Wfs1 knockout Mice at 7 Months of Age

To evaluate RGC loss, mouse retinal sections were stained with anti-Brn3a and anti-RBPMS, specific markers for RGC (Figure 2A). As previously reported, anti-RBPMS labels a greater number of RGCs than anti-Brn3a [34]. Results showed no evidence of RGC loss in the GCL of *Wfs1* knockout mice compared with wildtype male littermates using either Brn3a (Figure 2B) or RBPMS (Figure 2C) analysis at 7 months of age. These results indicate that loss of *Wfs1* does not induce RGC loss at 7 months of age, suggesting loss may occur at a more advanced stage in this knockout mouse model.

**Figure 2.**
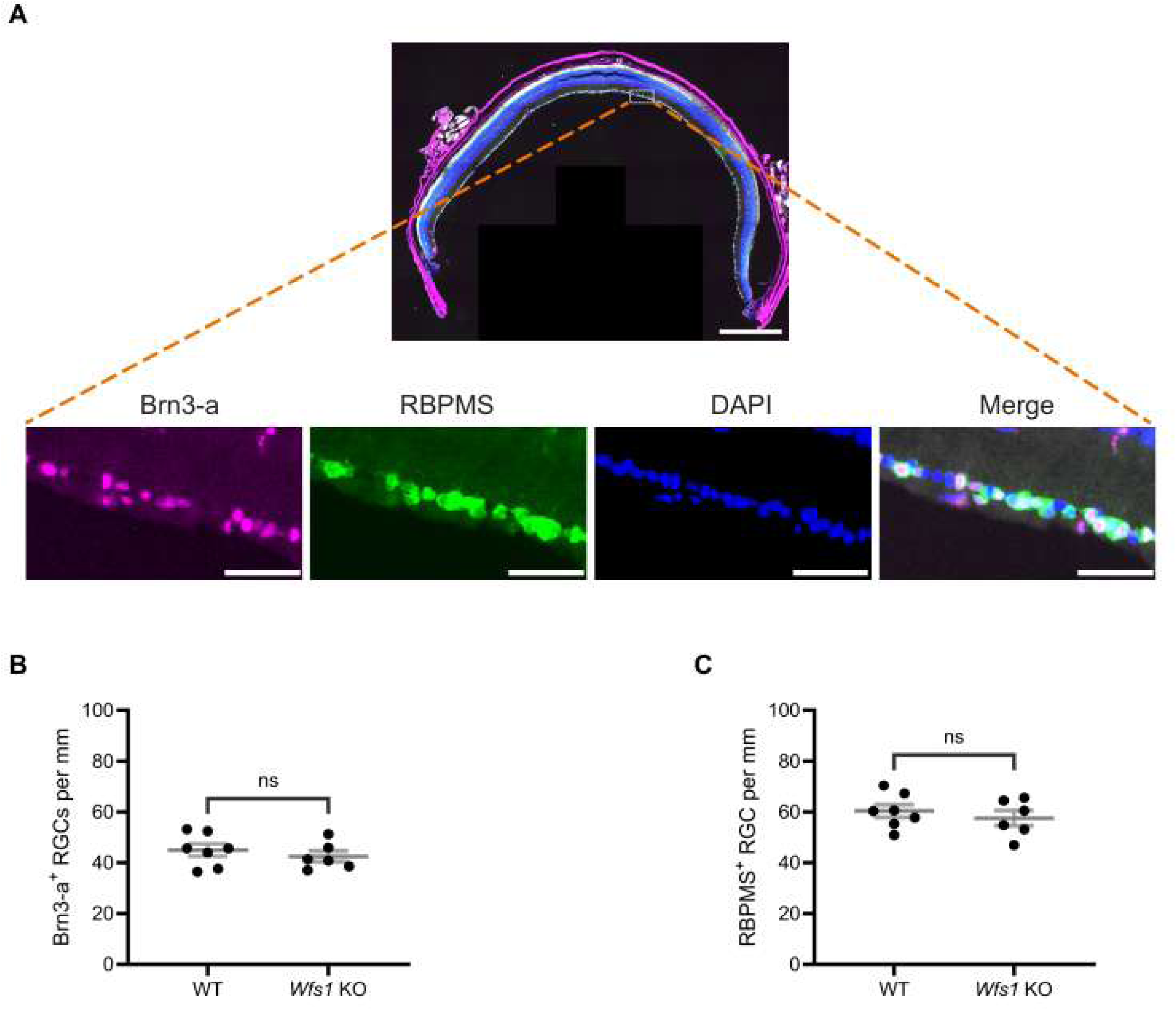
No RGC loss was evident in the *Wfs1-KO* model. (A) A representative mouse retinal section immunostained with both anti-Brn3-a (Magenta) and anti-RBPMS (green) antibodies counterstained with DAPI (blue). Scale bar. 500 µm. High-magnification insets of the boxed area show the individual fluorescence channel and the merged overlay with a scale bar of 50 µm. Quantification of (B) Brn3a-positive retinal ganglion cells (C) RBPMS-positive retinal ganglion cells in both *Wfs1* KO and control wild type mice of 7 month age. Quantification of results were presented as mean ± SEM. Statistical analysis was performed by two-tailed, unpaired student’s t-test with Welch’s correction. n ≥ 6. ns - not significant.

### 3.3 No Sign of Dendritic Loss in Wfs1 knockout Model

To evaluate dendritic loss in the inner plexiform layer (IPL), retinal sections from Wild type (WT) and *Wfs1* knockout mice at 4 and 7 months of age were immunolabeled with anti-β-III tubulin, a marker of neuronal soma, dendrites, and axons [33]. β-III tubulin labeling in the IPL showed no difference between WT and *Wfs1* knockout mice at either age (Figure 3A). Quantification of β-III tubulin mean intensity showed no significant difference at 4 months (*Wfs1* knockout: 0.9048 ± 0.046 vs. WT: 1.000 ± 0.059; Figure 3B) or at 7 months (*Wfs1* knockout: 1.022 ± 0.021 vs. WT: 1.000 ± 0.031; Figure 3C). A similar pattern was observed with β-III tubulin staining volume, with no significant changes in *Wfs1* knockout mice compared to age- and sex-matched littermate controls, indicating no dendritic loss in the IPL.

**Figure 3.**
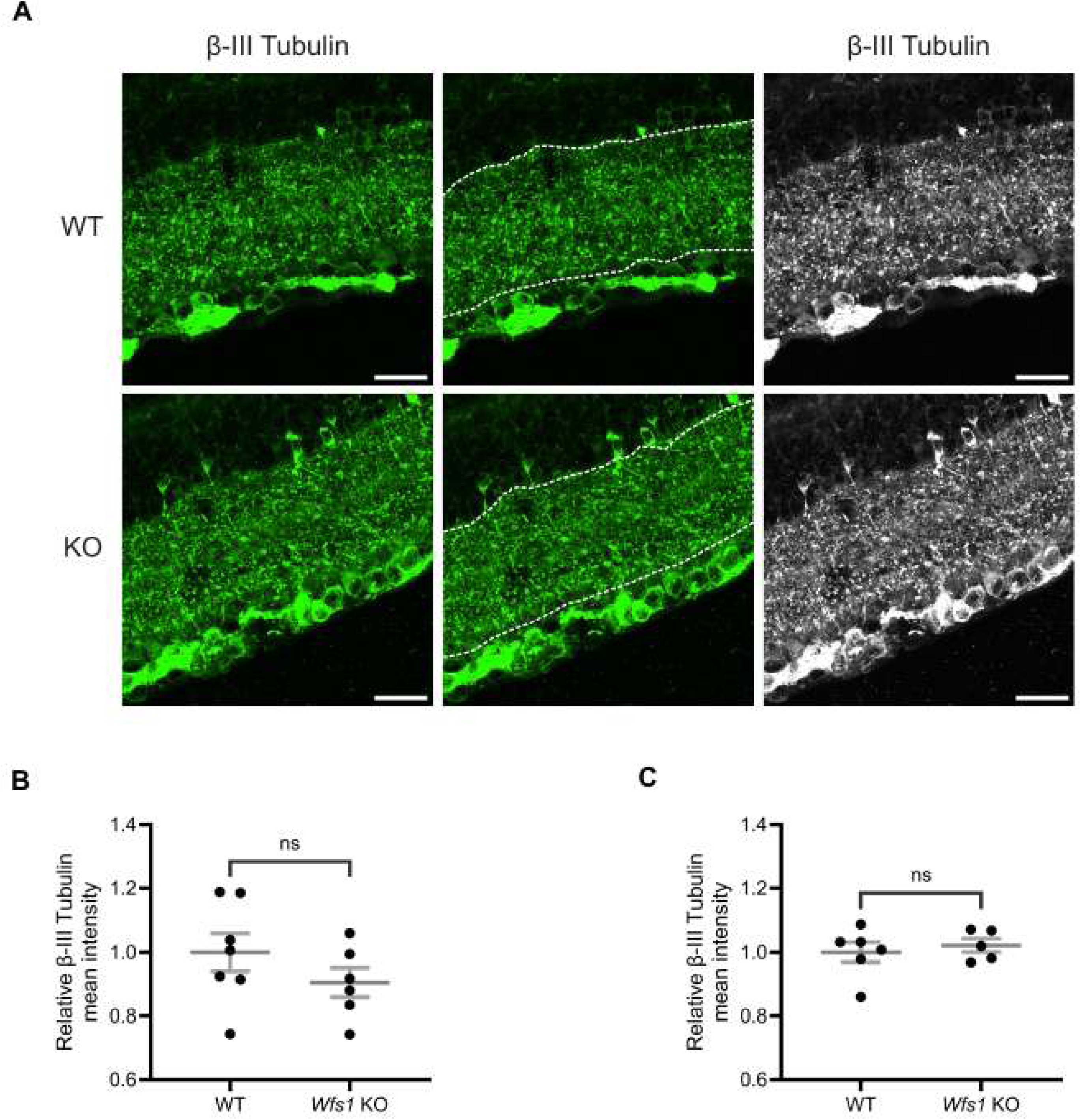
Dendritic architecture was preserved in *Wfs1* KO mice at 4 and 7 months. (A) Retinal sections were immunolabeled with an anti-13-111 tubulin antibody (green). Representative confocal images from 7-month-old WT (top) and *Wfs1* KO (bottom) retinas are shown. Scale bar, 20µm. Quantification of β-III tubulin fluorescence mean intensity in retinal sections from (B) 4-month and (C) 7-month old animals. Data are expressed as normalized values ± SEM relative to WT. Statistical comparisons were performed using a two-tailed unpaired Student’s t-test with Welch’s correction. n ≥4. ns - not significant.

### 3.4 Synaptic Alterations Are the Earliest Phenotype in Wfs1 knockout Model

To characterize the retinal phenotype further, we examined synaptic connections in the IPL, where RGCs form synapses through their dendrites with bipolar and amacrine cells (Yu et al., 2013). Retinal sections from WT and *Wfs1* knockout mice at 4 months (Figure 4A) and 7 months (Figure 5A) were immunostained with anti-PSD95 (postsynaptic density marker) and anti-synaptophysin (SYP; presynaptic marker).

**Figure 4.**
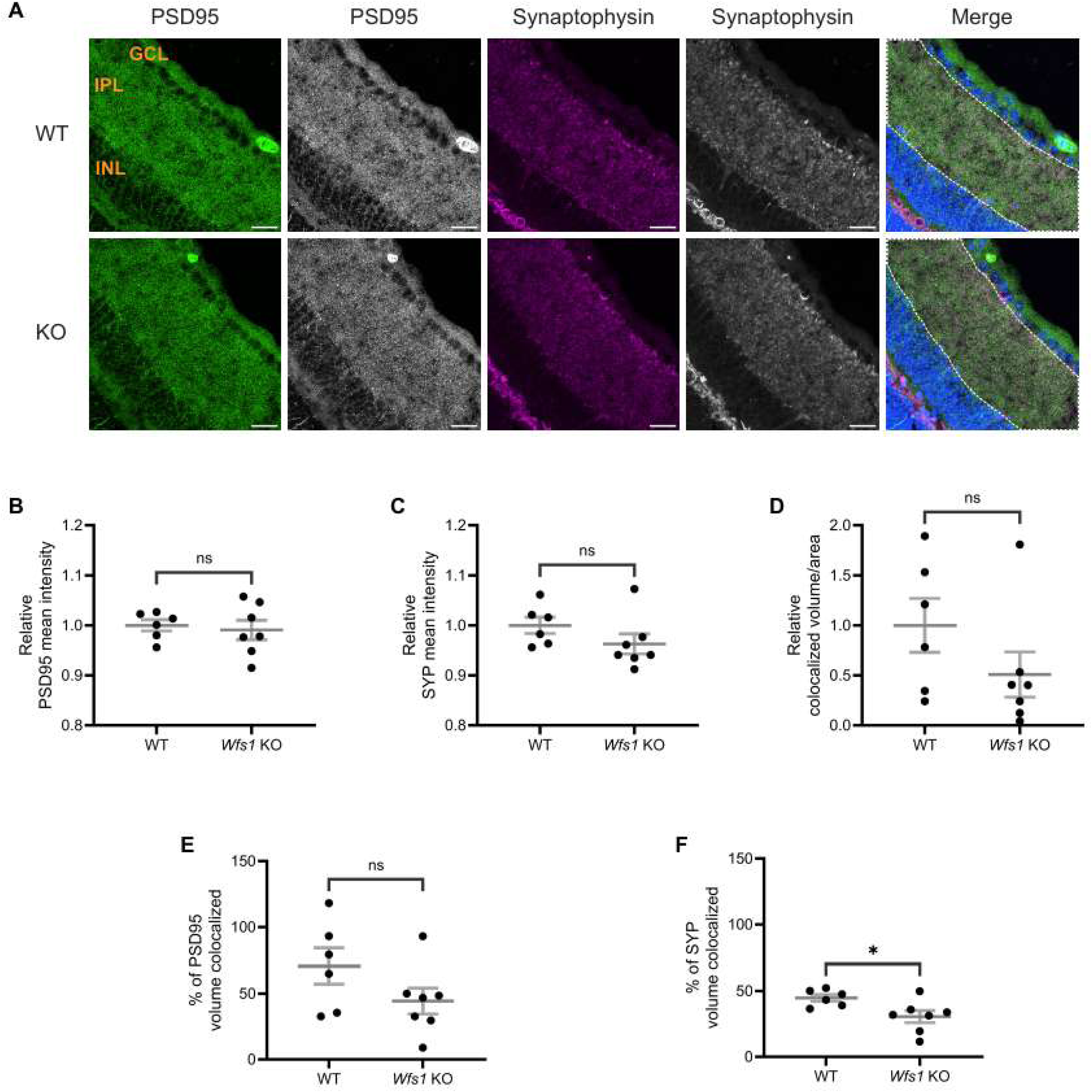
Altered synaptic connections were observed in *Wfs1* KO retinas at 4 months. (A) Retinal sections were immunolabeled with an anti-PSD95 antibody (green) and anti-synaptophysin antibody (Magenta). Representative confocal images from 4-month-old WT (top) and *Wfs1* KO (bottom) mice are shown. Scale bar, 20µm. Quantification of (B) PSD95 fluorescence mean intensity (C) synaptophysin fluorescence mean intensity in retinal sections from 4-month-old animals. Data are presented as normalized values ± SEM relative to WT. (D) Quantification of relative colocalized volume of PSD95-synaptophysin per unit ROI area in 4-month-old mice.(B) Quantification of relative percentage of (E) PSD95 (F) synaptophysin colocalized volume in 4 month old mice. Data are presented as normalized values ± SEM relative to WT. Statistical analysis was performed using a two-tailed unpaired Student’s I-test with Welch’s correction. n;,:6. *\*p* < 0.05, ns - not significant.

**Figure 5.**
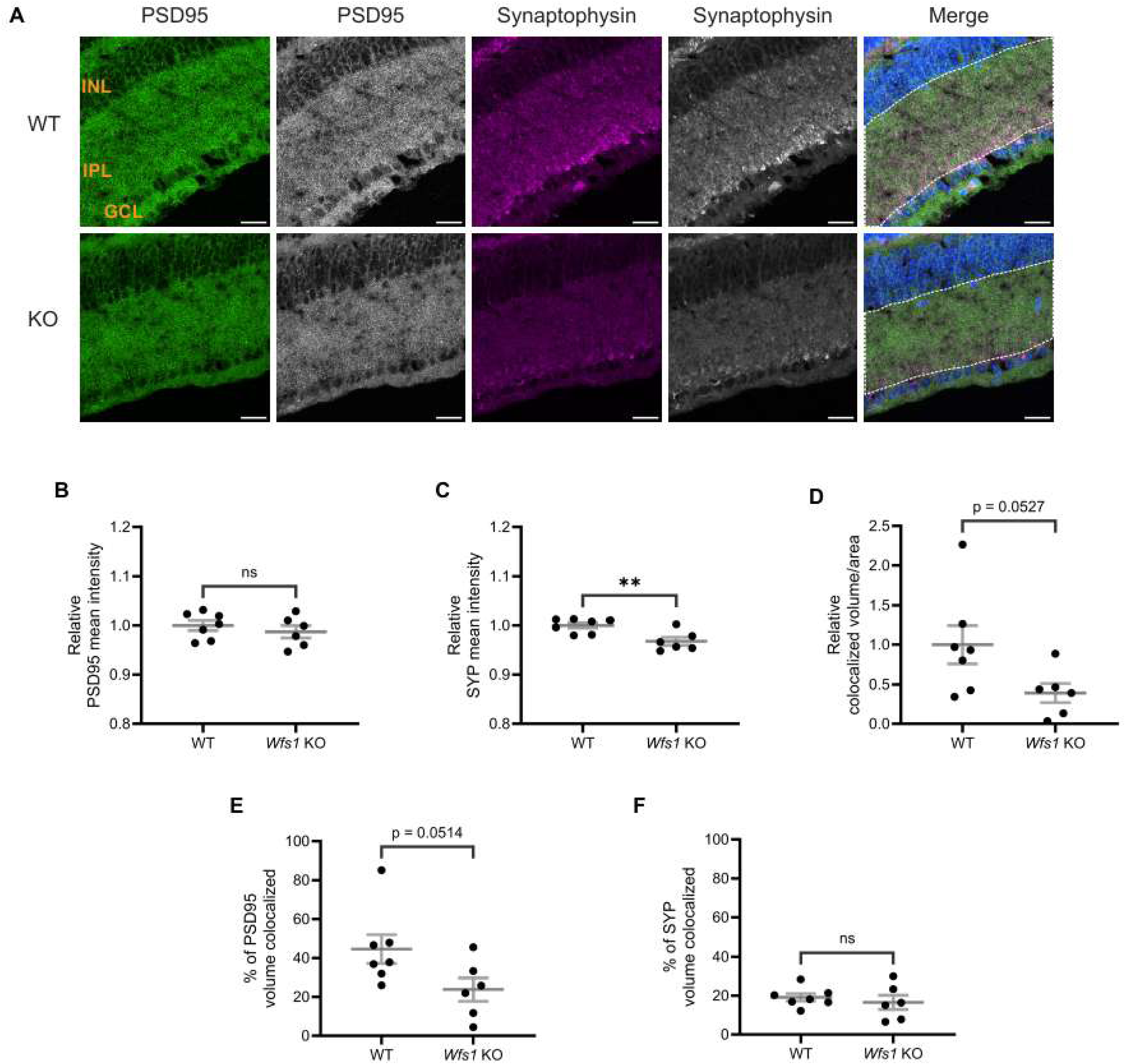
Synaptic loss was observed in *Wfs1* KO retinas at 7 months. (A) Retinal sections were immunolabeled with an anti-PSD95 antibody (green) and anti-synaptophysin antibody (Magenta). Representative confocal images from 7-month-old WT (top) and *Wfs1* KO (bottom) mice are shown. Scale bar, 20 µm. Quantification of (8) PSD95 fluorescence mean intensity (C) synaptophysin fluorescence mean intensity in retinal sections from 7-month-old animals. Data are presented as normalized values ± SEM relative to WT. (D} Quantification of relative colocalized volume of PSD95-synaptophysin per unit ROI area in 7-month-old mice.(B) Quantification of relative percentage of (E) PSD95 (F) synaptophysin colocalized volume in 7 month old mice. Data are presented as normalized values ± SEM relative to WT. Statistical analysis was performed using a two-tailed unpaired Student’s t-test with Welch’s correction. n ≥:6. *\*p* < 0.05, ns - not significant.

At 4 months of age, quantification of PSD95 mean intensity showed no difference (WT: 1.000 ± 0.011; vs. *Wfs1* knockout: 0.9910 ± 0.019; Figure 4B). SYP mean intensity was also similar between genotypes (WT: 1.000 ± 0.016; *Wfs1* knockout: 0.9627 ± 0.019; Figure 4C). Quantification of staining volume of PSD95 and SYP showed similar results (Supplementary Figure 1A-B). Object-based colocalization of PSD95 and SYP showed no statistically significant difference in colocalized volume per unit area (WT: 1.000 ± 0.269 vs. *Wfs1* knockout: 0.5081 ± 0.226; Figure 4D). However, the percentage of SYP volume colocalized with PSD95 was significantly decreased in *Wfs1* knockout mice (WT: 44.80 ± 2.557 vs. *Wfs1* knockout 30.56 ± 4.612 vs.; *p* = 0.0239; Figure 4F), while no significant difference was observed in the percentage of PSD95 volume colocalized (*Wfs1* knockout: 44.23 ± 9.802 vs. WT: 70.60 ± 13.60; Figure 4E).

At 7 months of age, PSD95 mean intensity remained unchanged (WT: 1.000 ± 0.010 vs. *Wfs1* KO: 0.9872 ± 0.012; Figure 5B). By contrast, SYP mean intensity was significantly decreased in *Wfs1* KO mice (WT: 1.000 ± 0.005 vs. Wfs1 knockout: 0.9677 ± 0.008; *p* = 0.009; Figure 5C). Similar results were observed with quantification analysis of staining volume of PSD95 and SYP (Supplementary Figure 1C-D). Colocalization analysis showed a 60% reduction in colocalized volume per unit area in *Wfs1* KO mice (0.3921 ± 0.121 vs. WT: 1.000 ± 0.242; *p* = 0.0527; Figure 5D). Quantification of the percentage of PSD95 volume colocalized revealed an approximately 50% reduction in *Wfs1* KO mice (23.80 ± 6.027 vs. WT: 44.60 ± 7.350; p = 0.0514; Figure 5E). The percentage of SYP volume colocalized showed no significant difference (Wfs1 KO: 16.55 ± 3.679 vs. WT: 19.16 ± 1.905; Figure 5F). Taken together, these results demonstrate that synaptic alterations are present at 4 months of age and progress to significant loss of synaptic connections by 7 months.

### 3.5 No Sign of Reactive Gliosis in Retina or Optic Nerve of Wfs1 KO Mice

To determine whether synaptic alterations were accompanied by glial activation, retinal sections were labeled with anti-GFAP. GFAP staining was largely restricted to the GCL and RNFL in both WT and *Wfs1* knockout mice at 7 months of age, indicating no extended glial processes (Figure 6A). Quantification of GFAP staining volume at 7 months showed a small but non-significant increase in *Wfs1* knockout mice (WT: 1.000 ± 0.154 vs. Wfs1 knockout: 1.281 ± 0.107; Figure 6C); the 4-month group showed comparable values (WT: 1.000 ± 0.091 vs. *Wfs1* knockout: 0.9003 ± 0.120; Figure 6B). A similar trend was observed for GFAP mean intensity (Supplementary Figure 2A–B).

**Figure 6.**
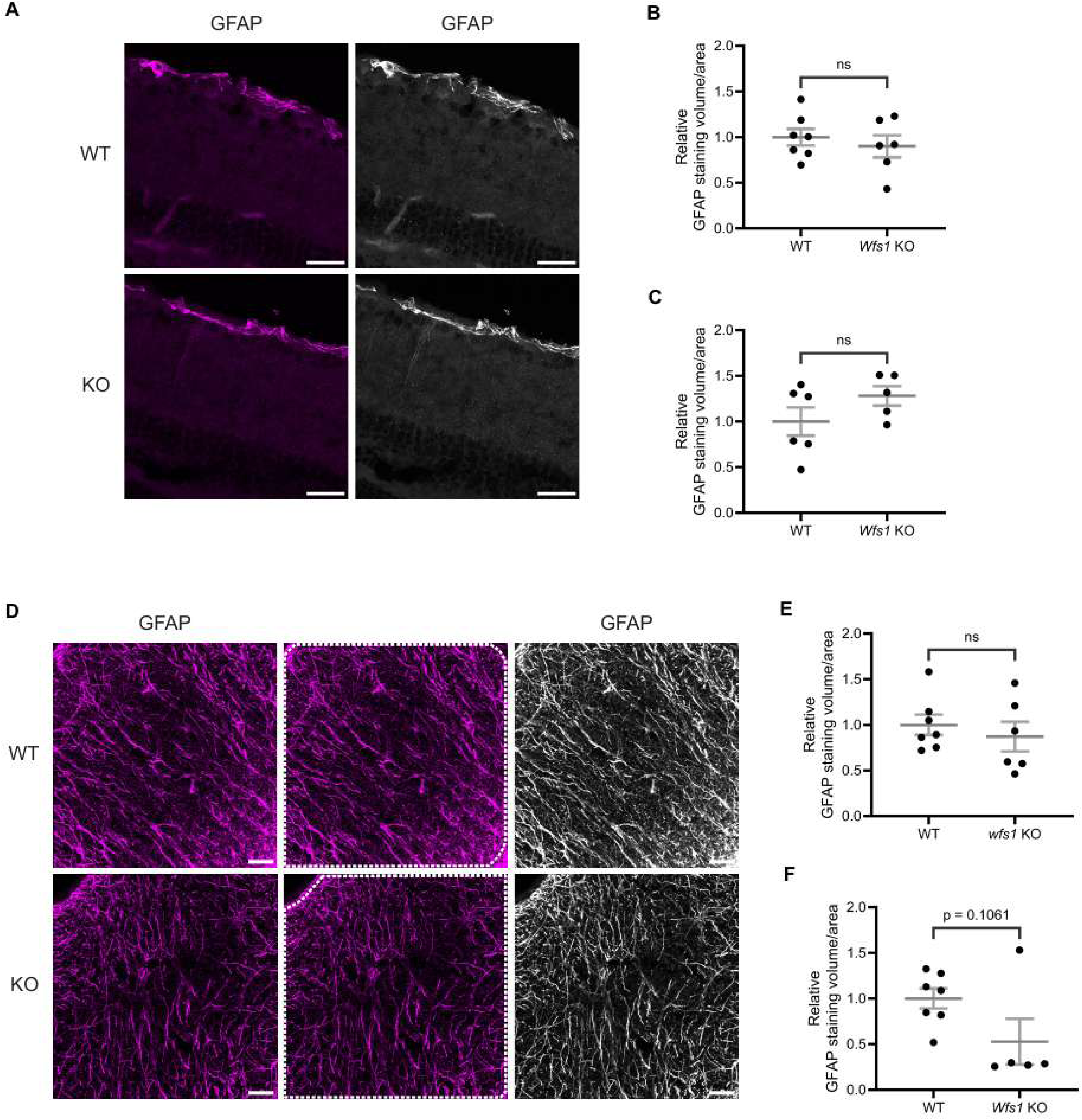
Elevated reactive gliosis was not evident in retina and optic nerve of *Wfs1* KO mice. (A) Retinal sections were immunostained with an anti-GFAP antibody (Magenta). Representative confocal images from 7-month-old WT (top) and *Wfs1* KO (bottom) retinas are shown. Scale bar, 20 µm. Quantification of relative GFAP-positive staining volume in retinal sections from (B) 4-month (C) 7-month-old mice. Values represent normalized means ± SEM relative to WT. Statistical analysis was performed using a two-tailed unpaired Student’s t-test with Welch’s correction. n ≥ 5. ns - not significant. (D) Optic nerve sections were immunostained with an anti-GFAP antibody (Magenta). Representative confocal images from 7-month-old WT (top) and Wfs1 KO (bottom) retinas are shown. Scale bar, 25 µm. Quantification of relative GFAP-positive staining volume in optic nerve sections from (E) 4-month (F) 7-month-old mice. Values represent normalized means ± SEM relative to WT. Statistical analysis was performed using a two-tailed unpaired Student’s I-test with Welch’s correction. n ≥ 5. ns - not significant.

Optic nerve sections from 4- and 7-month-old mice were also labeled with anti-GFAP. No sign of gliosis was observed in *Wfs1*-knokcout optic nerves at 7 months (Figure 6D). Strikingly, quantification of GFAP staining volume revealed an approximately 50% reduction in *Wfs1* knockout mice (WT: 1.000 ± 0.108 vs. Wfs1 knockout: 0.5276 ± 0.250; Figure 6F), while the 4-month group showed comparable values (WT: 1.000 ± 0.112 vs. *Wfs1* knockout: 0.8716 ± 0.162; Figure 6E). A similar trend was observed for GFAP mean intensity in optic nerves at both ages (Supplementary Figure 2C–D). Collectively, these results indicate an absence of reactive gliosis in both retina and optic nerve of *Wfs1* knockout mice at 4 and 7 months of age.

### 3.6 Significant Axonal Loss in the Optic Nerve of Wfs1 KO Mice at 7 Months

To evaluate axonal loss, a hallmark of optic atrophy, optic nerve transverse sections were immunolabeled with anti-neurofilament 200 (NF200) at 4 and 7 months of age. NF200 labeling was visibly reduced in *Wfs1* knockout animals at 7 months compared to wildtype controls (Figure 7A). Quantification of NF200 counts for the 7-month group yielded an average of 315,838 ± 28,571 per mm² in WT compared to 205,618 ± 46,408 per mm² in *Wfs1* knockout mice, a trend that approached significance (p = 0.0832; Figure 7D). Quantification of NF200 sum intensity per unit area showed a significant decrease in *Wfs1* knockout animals (WT: 1.000 ± 0.1698 vs. *Wfs1* knockout: 0.4428 ± 0.1778; p = 0.048; Figure 7E). In addition to this, quantification of the relative NF200-staining area fraction revealed a significant reduction, approximately 65%, in *Wfs1* knockout animals (WT: 1.00 ± 0.238 vs. *Wfs1* knockout: 0.35 ± 0.059; p = 0.033; Supplementary Figure 3). However, quantification of relative mean intensity showed reduction in *Wfs1* KO mice compared to WT mice but did not reach statistical significance (Data not shown).

**Figure 7.**
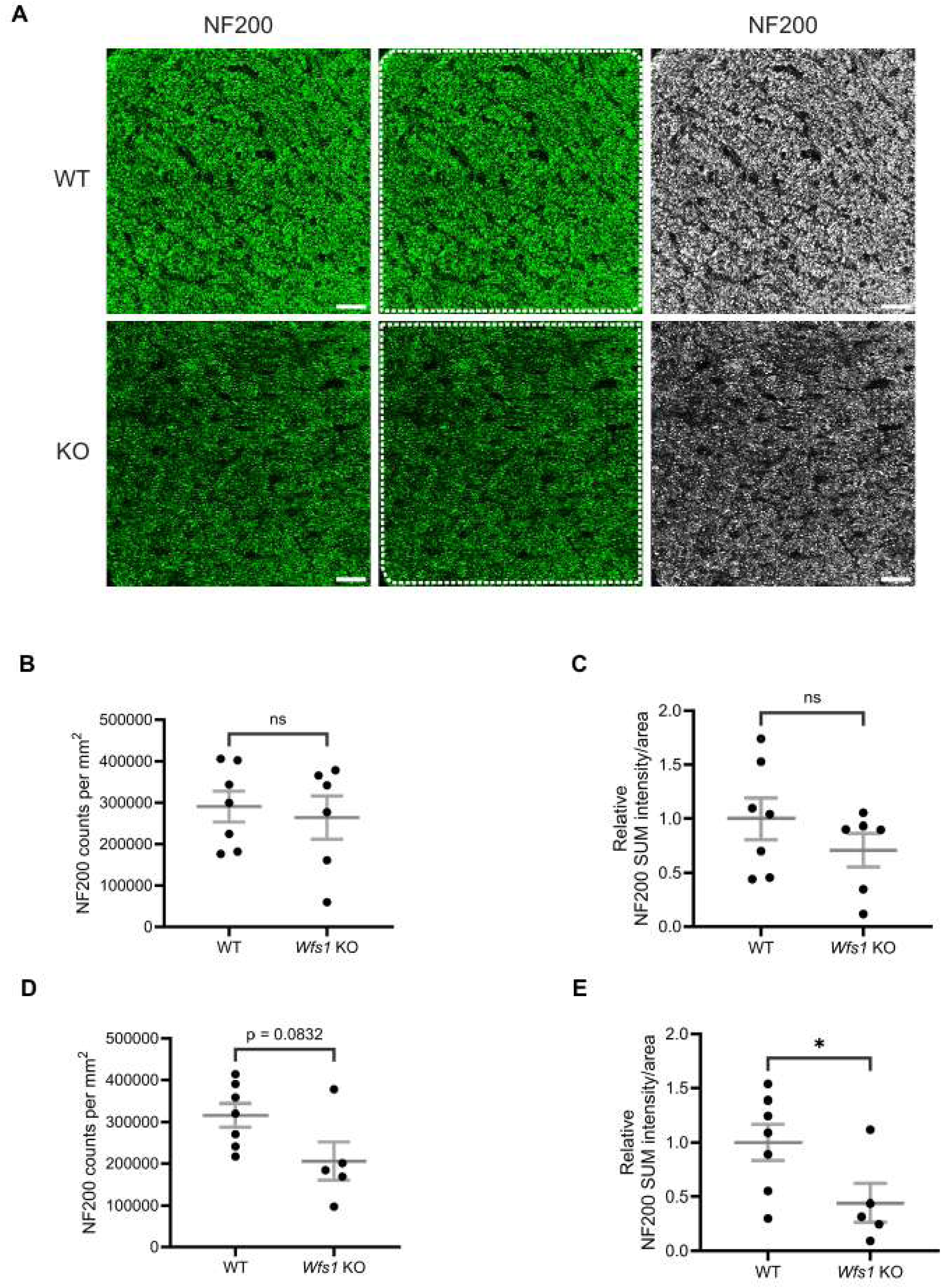
Axonal loss was observed in *Wfs1* KO mouse model. (A) Mouse optic nerve sections were immunolabeled with an anti-NF200 antibody (green). Representative confocal images obtained for 7-month-old WT (top) and *Wfs1* KO (bottom) mice are shown. Scale bar, 25 µm. Quantification of (B) axonal counts per mm^2^ and (C) NF200 staining volume per unit ROI area in the optic nerve sections of 4 month old mice. Quantification of (D) axonal counts per mm^2^ and (E) NF200 staining volume per unit ROI area in the optic nerve sections of 7 month old mice. Axonal counts were presented as mean ± SEM. NF200 staining volume results were presented as normalized values± SEM to the mean of WT group. Statistical analysis was performed by two-tailed, unpaired student’s t-test with welch’s correction. n ≥ 5. *P<0.05, ns - not significant.

In the 4-month group, NF200 counts were comparable (WT: 290,577 ± 37,146; *Wfs1* knockout: 263,909 ± 52,276; Figure 7B), and NF200 sum intensity showed only a modest, non-significant reduction (WT: 1.000 ± 0.1910; *Wfs1* knockout: 0.7090 ± 0.1553) (Figure 7C). These results demonstrate progressive, age-dependent axonal loss that is significant at 7 months in *Wfs1* knockout mice.

### 3.7 No Sign of Demyelination in Wfs1 knockout Model

To determine whether axonal loss was accompanied by demyelination, optic nerve sections were immunostained with anti-myelin basic protein (MBP) at 4 and 7 months of age. MBP staining showed no difference in myelination between WT and *Wfs1* knockout animals at either age (Figure 8A). Quantification of MBP mean intensity confirmed no significant difference at 4 months (WT: 1.000 ± 0.037; *Wfs1* KO: 1.052 ± 0.056) (Figure 8B) or 7 months (WT: 1.000 ± 0.056; *Wfs1* knockout: 1.024 ± 0.105) (Figure 8C). These results clearly indicate an absence of demyelination in the *Wfs1* knockout mouse model even with age progression.

**Figure 8.**
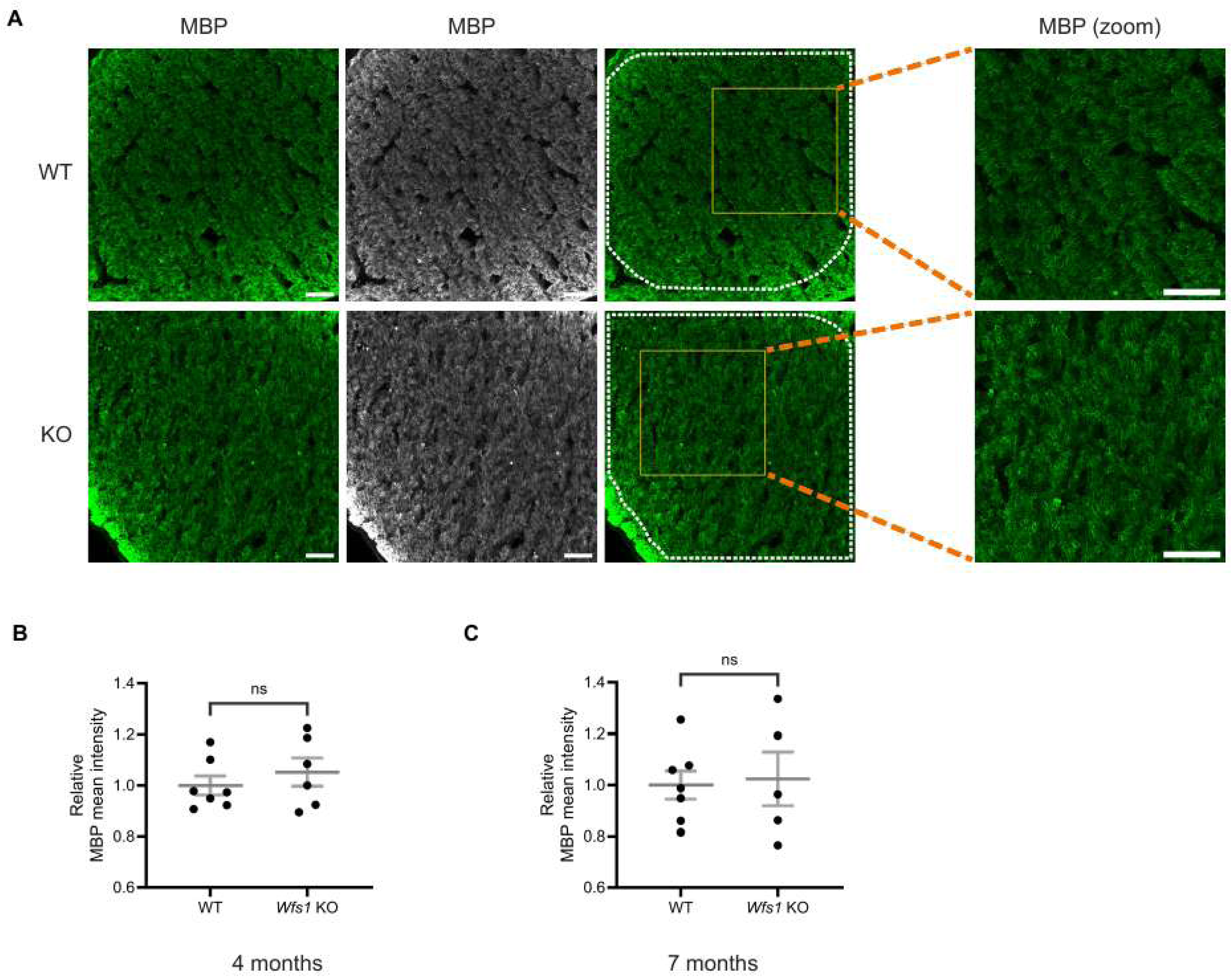
No evidence of demyelination was detected in optic nerves of *Wfs1* KO mice at 4 and 7 months. (A) Optic nerve sections were immunostained with an anti-MBP antibody (green). Representative confocal images from 7-month-old WT (top) and *Wfs1* KO (bottom) mice are shown along with the corresponding high-magnification zoom of the boxed area. Scale bar, 50 µm. Quantification of MBP fluorescence mean intensity in optic nerve sections from (B) 4-month (C) 7-month old animals. Data are presented as normalized values ± SEM relative to the WT mean. Statistical analysis was performed using a two-tailed unpaired Student’s I-test with Welch’s correction. n≥ 5. ns - not significant.

**Figure 9.**
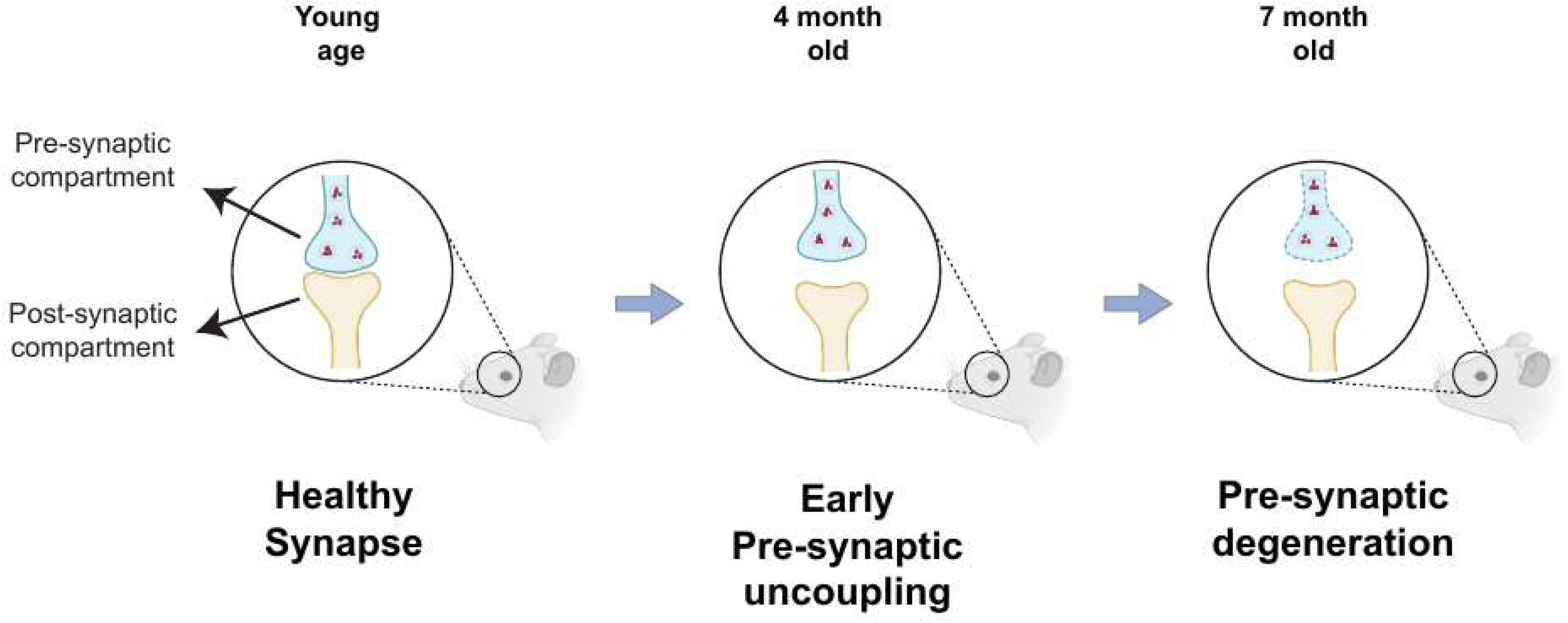
Schematic model of progressive synaptic loss in retina of Wfs1 KO mouse model. At younger age, retina of KO mouse show a intact and healthy synapse with pre- and post-synaptic compartments apposing each other closely in IPL of retina. At 4 months of age, pre-synaptic compartment showed early stage of uncoupling or spatial de-alignment which was evident with significant decrease of percentage of synaptophysin volume colocalized but not that of PSD95. At 7 months of age, pre-synaptic degeneration was evident with significant decrease of synaptophysin levels and overall decrease in colocalized volume with PSD95. This loss of synapse was accompanied with axonal loss at 7 months of age.

## 4. Discussion

Wolfram syndrome is a rare genetic disease characterized by early-onset antibody-negative diabetes mellitus, optic atrophy, and neurodegeneration. Preclinical models in mouse, rat, and zebrafish have been developed to model Wolfram syndrome and have been shown to manifest visual defects including RGC loss, axonal loss, myelin degeneration, and reactive gliosis [21; 22; 23; 24; 25; 26; 27; 28]. However, no previous studies in these models investigated synaptic and dendritic connections. *In vitro* evidence from cerebral organoids and iPSC-derived neurons indicated that *WFS1*-deficient neurons exhibit disrupted synapses and altered neurites [29; 30], underscoring the need for *in vivo* investigation. The present study, for the first time, reveals that synaptic alterations precede axonal loss and represent the earliest phenotype in optic atrophy of the *Wfs1* knockout mouse model.

The IPL of the mammalian retina harbors the synapses and dendrites of RGCs, which form connections with bipolar and amacrine cells from the INL. Our study reveals that the percentage of SYP colocalized with PSD95 is significantly decreased in *Wfs1* knockout mice compared to WT at 4 months of age, despite no significant changes in the levels of pre- or postsynaptic compartments at that age, likely reflecting uncoupling of presynaptic compartments. As the disease progresses to 7 months, mutant mice exhibit loss of presynaptic compartments (decreased SYP mean intensity), reduced colocalized volume overall, and decreased percentage of PSD95 colocalized with SYP, indicating progressive loss of functional synapses. This pattern of early presynaptic uncoupling followed by eventual compartment loss at advanced disease stages indicates a key role for presynaptic failure in the mechanism of Wolfram syndrome. Notably, similar results were reported in cerebral organoids where *WFS1*-deficient neurons showed decreased colocalization of Synapsin 1 with PSD95 and reduced synapse density [29]. This early synaptic dysfunction may explain the visual defects observed well before axonal loss in both mouse and rat models [24; 27]. Furthermore, presynaptic failure is well documented as an early event in Alzheimer’s disease and dominant optic atrophy [35; 36; 37; 38; 39], while reduction of PSD95 has been shown to occur at a later stage in Alzheimer’s disease [40], consistent with the relatively stable postsynaptic compartment observed here in early-stage disease.

Further retinal investigation confirms no RGC loss and no dendritic loss in *Wfs1* knockout mice in this study. Using both Brn3a (which labels approximately 85% of murine RGCs)[41] and RBPMS (which labels nearly all RGCs)[34], we confirm no RGC loss at 7 months of age. We show no dendritic loss in the IPL at either 4 or 7 months, which contrasts with findings from human iPSC-derived neurons where neurite outgrowth was impaired [30]. This discrepancy may direct attention to sublamina-specific dendritic degeneration within the IPL, as has been described in dominant optic atrophy [42] and glaucoma [43].

In this study, we have not shown evidence any significant reactive gliosis in both retina and optic nerve. In fact, we observe almost 50% reduction of GFAP staining volume in the optic nerve of *Wfs1* knockout mice compared to the WT mice at 7 months of age whereas the retina shows a mild, non-significant increase in GFAP staining volume in the knockout animals. This anatomical discrepancy can be attributed to the heterogeneous distribution of macroglia; while the retina housing both Muller cells and astrocytes, the optic nerve is completely devoid of Muller cells but astrocytes. While no previous studies investigated reactive gliosis state in optic nerve, few studies revealed gliosis state in the retina of *Wfs1* knockout mouse model. Increased gliosis in the retina in *Wfs1* knockout mice was shown before [23]. This contradicting result can be attributed to the genetic background of the mice which was shown to impact the severity of the phenotype [36]. However, our findings are consistent with another study that showed reduced GFAP intensity in the retina of *Wfs1* knockout mice [22]. Furthermore, it was demonstrated that *WFS1*-deficient cerebral organoids had reduced number of astrocytes [29]. Altogether, our finding of reduced gliosis in the optic nerve indicates impaired astrocytic activity in *Wfs1* knockout mice and warrants further investigation to determine whether this change is a cause or a consequence of disease progression.

Regarding axonal integrity, we observe an approximately 40% reduction in NF200 intensity in *Wfs1* knockout mice compared to WT at 7 months, with no significant loss at 4 months. This result is consistent with previous reports of axonal loss without concurrent RGC degeneration [23], while the later onset relative to that study may be attributed to differences in genetic background [36]. No demyelination is detected at either age, though we cannot exclude subtle abnormalities that would require transmission electron microscopy (TEM) analysis to detect.

This study, for the first time, reveals synaptic alterations as the earliest disease phenotype in the optic atrophy of Wolfram syndrome, preceding axonal loss in *Wfs1* knockout mice. As the disease progresses, presynaptic failure leads to an overall loss of functional synapses. The involvement of presynaptic pathology in Alzheimer’s disease and dominant optic atrophy supports the relevance of investigating presynaptic machinery in Wolfram syndrome. Targeting early synaptic alterations may represent a promising therapeutic strategy at the pre-degenerative stage. In addition, this study identifies reduced astrocytic reactivity in the optic nerve as a novel phenotype, raising new questions about the role of astrocyte failure in Wolfram syndrome pathogenesis and progression.

## Author Contributions

VG and FU designed the study. VG. performed the experiments and analyzed the data. WA and SB assisted with experiments. VG and FU wrote the manuscript. All authors reviewed and approved the final version.

## Data Availability Statement

The raw data supporting the conclusions of this article will be made available by the authors, without undue reservation.

## Acknowledgments

The authors thank Dr. Sulev Kõks (University of Tartu) for providing the 129S6 Wfs1 KO mouse line. We also thank the Washington University School of Medicine imaging core facilities for their support and the members of the Washington University Wolfram Syndrome and Related Disorders Clinic and Research Team (https://wolframsyndrome.wustl.edu) for their invaluable support. We extend special thanks to Mr. Gabriel Skinner and Mr. Cris Brons for managing the mouse colony. We also gratefully acknowledge the philanthropic support and encouragement from the Auerbach Hyman Fund, the Philipp Fund, Jerome W. Gratenstein Memorial Foundation, the WAVE fund, the WAV fund, the Silberman Fund, the Stowe Fund, the Feiock Fund, the Cachia Fund, the Gildenhorn Fund, the Snow Foundation, the Ellie White Foundation for the Rare Genetic Disorders, the Unravel Wolfram Syndrome Fund, Be A Tiger Foundation, the Eye Hope Foundation, Ontario Wolfram League, Associazione Gentian Sindrome di Wolfram Italia, Alianza de Familias Afectadas por el Sindrome Wolfram Spain, Wolfram Heroes Society, Wolfram syndrome UK, and Association Syndrome de Wolfram France.

---

**Supplementary Figure 1.**
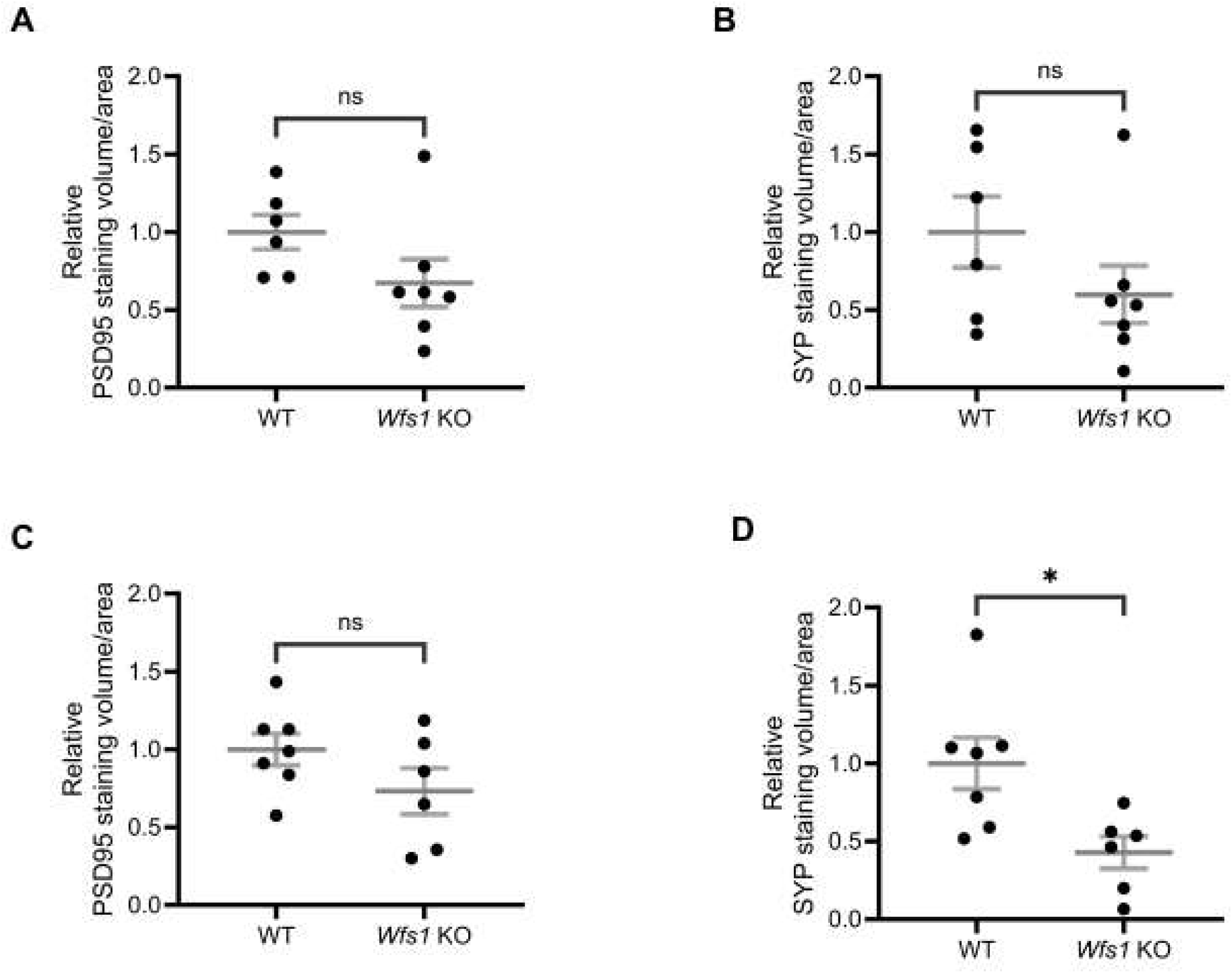
Quantification of synaptic proteins in *Wfs1* KO mice. Relative quantification of (A,C) PSD95 and (B,D) synaptophysin staining volume in 4 (A and B) and 7 (C and D) month old *Wfs1* KO mice compared to WT mice. Data are presented as normalized values± SEM relative to WT. Statistical comparisons were performed using a two-tailed unpaired Student’s t-test with Welch’s correction. n ≥ 6. *\*p* < 0.05. ns - not significant.

**Supplementary Figure 2.**
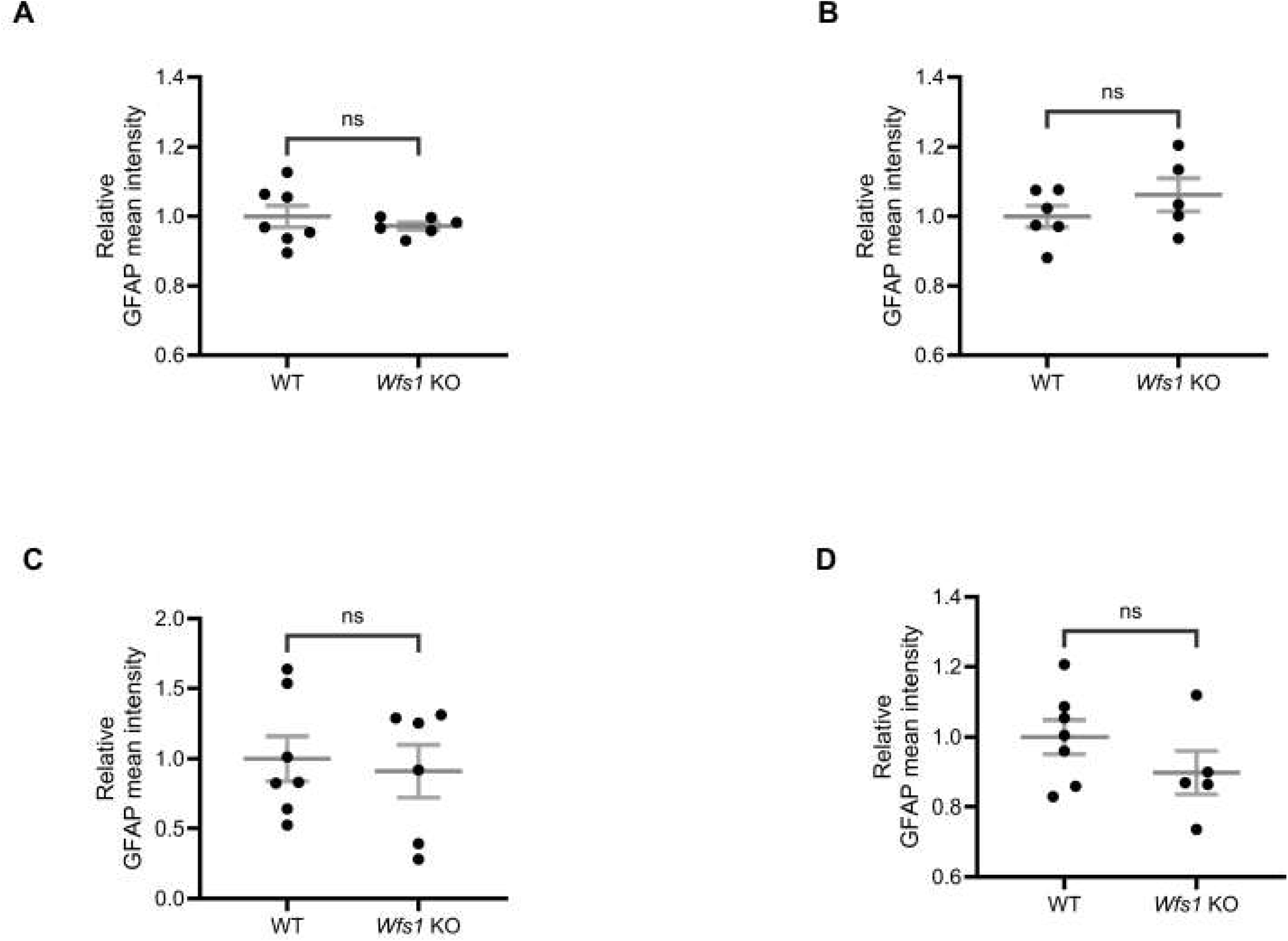
Quantification gliosis state in retina and optic nerve of *Wfs1-KO* mouse model. Relative quantification of GFAP staining mean intensity in retina (A and 8) and optic nerve (C and 0) of 4 (A and C) and 7 (8 and D) month old *Wfs1* KO mice compared to WT mice. Values represent normalized means ± SEM relative to WT. Statistical comparisons were performed using a two-tailed unpaired Student’s I-test with Welch’s correction. n ≥ 5. ns - not significant.

**Supplementary Figure 3.**
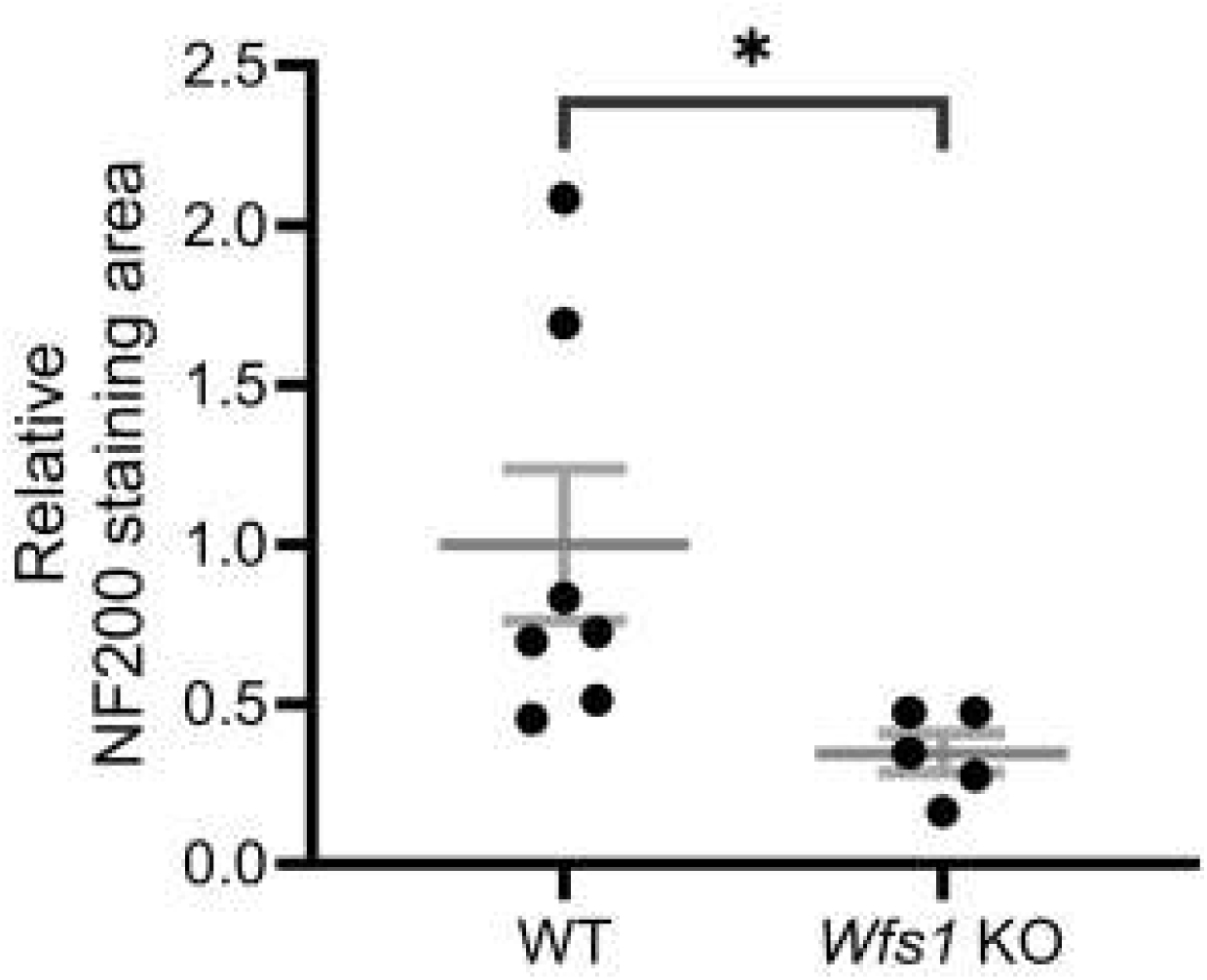
Quantification of NF200 staining area in optic nerve section of *Wfs1* KO model. Relative quantification of NF200 staining area in optic nerve sections of 7 month old *Wfs1* KO mice compared to WT mice. Data points represent normalized values ± SEM relative to WT. Statistical comparisons were performed using a two-tailed unpaired Student’s I-test with Welch’s correction. n ≥ 5. \**P*<0.05.

